# Real-time volatilomics reveals microbiota and pathogen fingerprints in the honey bee

**DOI:** 10.1101/2025.10.30.685588

**Authors:** Mateusz Fido, Silvia Moriano-Gutierrez, Jiayi Lan, Aurélien Pirat, Léa Zufferey, Elisa Cappio Barazzone, Renato Zenobi, Philip Engel, Emma Slack

**Author notes:** Joint first authors (Mateusz Fido, Silvia Moriano-Gutierrez).

## Abstract

Understanding the complex relationship between gut microbiota and their hosts often relies on invasive sampling techniques. Honey bees provide a tractable model for host-microbe studies. Here we establish single-bee volatilomics using secondary electrospray ionization (SESI-HRMS) to examine volatile organic compounds released to the air around an individual live honey bee. Specifically, we focused on primary gut microbiota metabolites present in gnotobiotic bees. Our findings reveal distinct volatilome profiles in honey bees that depend on their gut bacterial colonization state. We cross-validated our findings using an established metabolomics technique, LC-HRMS, to compare and contrast the metabolites detectable with each mass spectrometry-based method. Finally, we assessed the ability of SESI-HRMS to detect colonization with the bee pathogen *S. marcescens*. By comparing the volatile signature of this bacterium grown in liquid culture with that of infected honey bee headspace, we identified overlapping compounds, including butane-2,3-diol, that were absent in uninfected bees. SESI-HRMS volatilomics, paired with LC-HRMS, therefore has potential to identify non-invasive biomarkers of bee microbiome composition and infection at the level of individual insects. These biomarkers represent practical targets for the development of simple, field-ready diagnostic tools for monitoring pollinator health.

## INTRODUCTION

The gut microbiota is a critical component of host physiology, influencing a wide range of biological functions, including digestion, immune regulation, metabolism, and even behavior.^1–6^ This symbiotic association is mediated by a diverse array of biochemical interactions, many of which involve the production and exchange of small molecules. Among these, volatile organic compounds (VOCs) represent an important class of metabolites that can serve as indicators of both host physiology and microbial activity.^7,8^ VOCs are produced as metabolic byproducts by both the host and its associated microbiota and provide a unique window into the functional state of an organism.^9,10^ These compounds include alcohols, aldehydes, ketones, organic acids, and short-chain fatty acids (SCFAs), many of which are products of microbial fermentation.^1,11,12^ Beyond their metabolic roles, VOCs function as infochemicals in both intra- and interspecies communication and have been shown to influence host behavior, immune signaling, and disease states.^13,14^

Despite their biological importance, capturing these dynamic metabolic exchanges in a living system has traditionally been challenging. Conventional metabolomics requires invasive sampling techniques such as tissue dissection and offers only static snapshots of metabolic activity. Fecal sampling presents a less invasive alternative, though it can be temporally disconnected from systemic metabolism, reflecting only downstream processing and dominated by the microbial output. Recent advances in volatilomics have paved the way for non-invasive monitoring of metabolic processes in living organisms.^15–17^ Unlike traditional methods that require disruption of biological systems, volatilomics enables real-time assessment of molecular interactions by detecting VOCs released into an organism’s immediate environment, including low-abundance aerosolized compounds. While this approach has been successfully applied in medical diagnostics,^18^ food safety,^19^ and environmental monitoring,^20^ it remains underutilized in host-microbe interactions research.

Secondary electrospray ionization-high-resolution mass spectrometry (SESI-HRMS) is a novel analytical technique that enables the detection of VOCs in real time, without the need for sample preparation or manipulation.^21^ This approach utilizes an ambient ionization method to detect VOCs present in the headspace of a sample, allowing for non-invasive and dynamic monitoring of host and microbial metabolism. The SESI-HRMS technique leverages the principles of electrospray ionization to detect VOCs present in the headspace of honey bee samples. In this method, a gentle flow of a carrier gas directs VOCs towards an electrospray plume, where they are subsequently ionized by charged species in the electrospray and separated based on their mass-to-charge ratio in the mass analyzer. This strategy benefits from very high sensitivity and selectivity of high-resolution mass spectrometry compared to conventional mass spectrometry methods, allowing for the discrimination of VOCs with similar molecular weights but distinct chemical structures.

The Western honey bee (*Apis mellifera*) serves as an excellent model for investigating host-microbiota interactions through volatilomic profiling. It harbors a relatively simple and well-characterized gut microbiota,^22–26^ and represents an experimentally tractable system that allows to colonize microbiota depleted bees with cultured bacteria assembled into defined communities.^27–29^ This allows for controlled studies on microbial contributions to host metabolism and volatile production. In honey bees, VOCs play essential roles in chemical communication, foraging, and colony organization.^30–32^ Recent studies have also shown that the gut microbiota contributes directly to VOC production, including short-chain fatty acids (SCFAs) and other fermentation-derived metabolites that shape colony odor and host signaling.^33,28,34^

Despite their ecological and economic importance, honey bee populations are experiencing global declines driven by a combination of stressors, including habitat loss, pesticide exposure, nutritional stress, and pathogenic infections.^35,36^ Opportunistic pathogens such as *Serratia marcescens* are emerging as significant contributors to honey bee mortality, particularly in individuals with disturbed or depleted microbiota.^37^ Current methods for detecting infections or monitoring bee health are limited in resolution, invasiveness, or scalability. Therefore, there is a pressing need for non-invasive, real-time diagnostic tools capable of detecting microbial dysbiosis or pathogen colonization at the individual level.

In this study, we utilize SESI-HRMS to non-invasively characterize the volatilome of living honey bees. By comparing microbiota-depleted bees with those colonized by a synthetic microbial community, we aim to disentangle the contributions of host and microbial metabolism to VOC production and shed light on the critical role of gut microbiota in shaping the chemical signatures of honey bees. This study not only provides new insights into the metabolic interactions between gut microbiota and their insect hosts, contributing to a deeper understanding of microbiome function, but also facilitates the identification of VOC biomarkers linked to microbiota activity. In addition to exploring the role of core gut microbes, this study extends the application of SESI-HRMS to the *in vivo* detection of pathogenic bacteria, specifically *Serratia marcescens*, an opportunistic pathogen increasingly implicated in honey bee mortality and colony collapse.^38^ We identify and characterize the VOC signatures associated with *Serratia* colonization, both in the bee gut and in the headspace of living individuals. These biomarkers could pave the way for novel, non-invasive diagnostic tools for monitoring honey bee health. Given the global decline in pollinator populations, microbiota-based indicators of bee health could play a crucial role in informing beekeeping strategies and improving colony management practices.

## RESULTS

### Gut microbiota colonization drives distinct volatilome metabolic profiles in honey bees

To investigate how gut microbiota shape VOC emissions in honey bees, we compared the headspace profiles of microbiota-depleted (MD) bees and those colonized with a defined synthetic community (SC) using single-bee SESI-HRMS (Fig. 1A). Bees were drawn from four independent hives (1-4). Within each hive, individuals were housed in two separate cages (A and B) to account for potential cage effects, and two bees per cage were measured independently by single-bee SESI-HRMS (16 total single-bee measurements).

**Fig 1.**
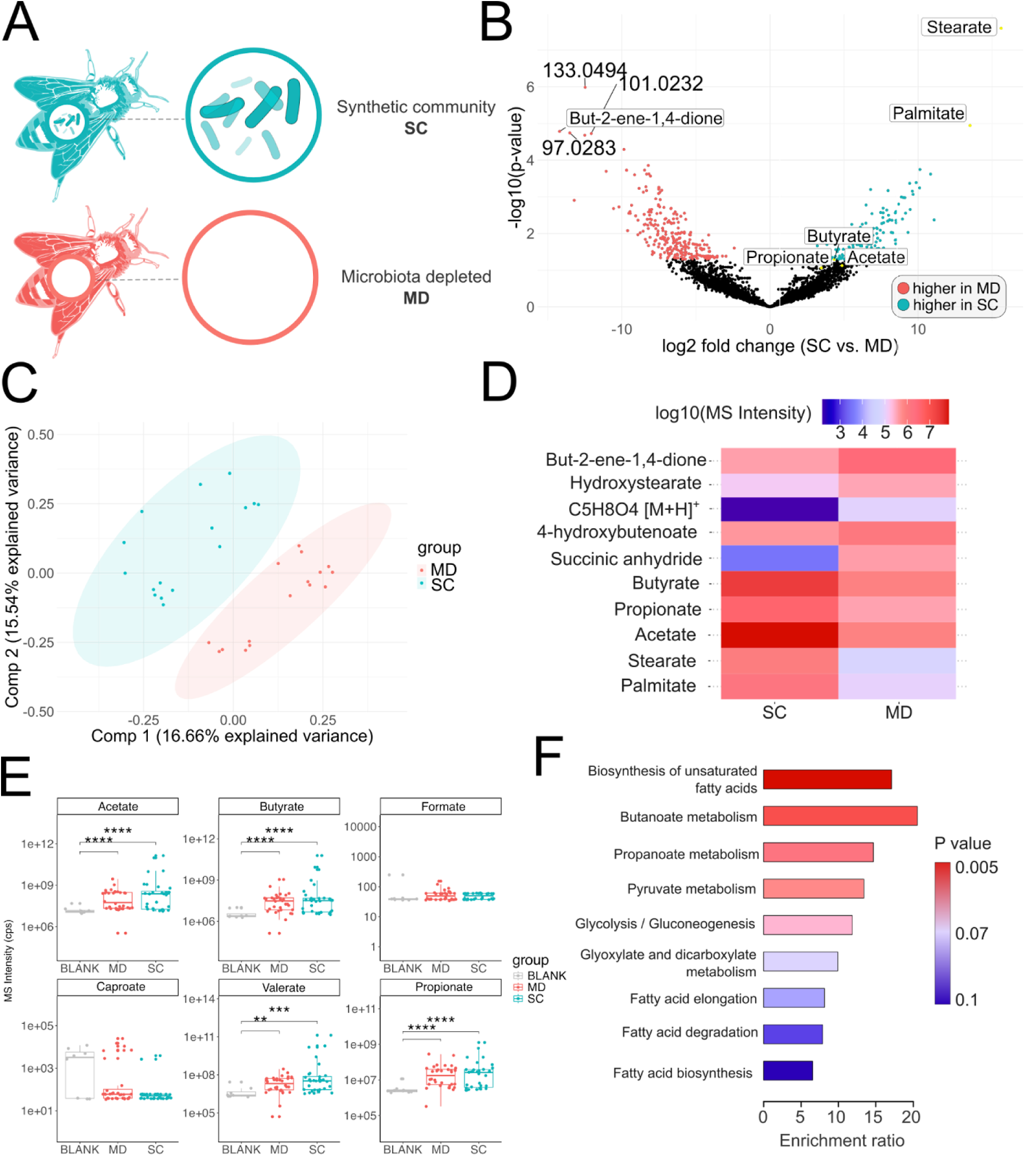
Single-bee measurements of honey bee headspace using SESI-HRMS. **A**) Schematic of the experimental design. **B)** Volcano plot showing the differential abundances of features shared between the SC and MD groups. Features with p < 0.05 and fold change greater than 2 are highlighted in blue (higher in SC) and red (higher in MD). **C)** PLS-DA plot showing the sample distribution between the SC and MD groups. **D)** Heatmap showing the logarithmic MS intensity of the most significantly different compounds in SC and MD bee guts by SESI-HRMS. The intensities correspond to MS/MS-confirmed m/z values of identified metabolites. **E)** Mean ion counts corresponding to short-chain fatty acids, matched by accurate mass. Statistical significance was measured by Welch’s t-test, adjusted for multiple comparisons using the Holm-Bonferroni method. P-value cutoff points: * - p < 0.05, ** - p < 0.01, *** - p < 0.001, **** - p < 0.0001. Blank samples correspond to measurements of an empty PMMA tube connected to the carrier gas but without a bee inside. **F)** Enrichment analysis of the differentially abundant MS/MS-identified compounds from panel D (MetaboAnalyst 5.0, KEGG metabolite set).

To check the gnotobiotic status of the sample bees, we carried out qPCR with universal bacterial primers and full-length 16S rRNA amplicon sequencing (Supporting Information, Fig. S1-S3). These analyses confirmed that bacterial loads were much lower in MD bees as compared to SC bees and that the SC bees were colonized by the expected bacteria while MD bees contained several non-specific strains, possibly coming from contamination or PCR artifacts (SI, Fig. S1).

The gut microbiota colonization status profoundly impacted the global volatile profile of the live animal. Out of 3570 detected features, 420 showed differential abundance m/z values between SC and MD bees, with 259 enriched in MD bees and 161 in SC bees (Fig 1B). A clear separation between the two groups was observed in the PLS-DA score plot (Fig. 1C), providing more evidence that microbial colonization drives distinct volatile metabolic signatures at the individual level.

From the differential abundance analysis and PLS-DA loadings, nine key metabolites were identified by MS/MS (Fig. 1D). These included several SCFAs and related intermediates such as acetate, propionate, butyrate and succinic anhydride, all canonical end products of microbial fermentation. Despite low abundance, acetate, propionate, and butyrate all showed above-background ion counts (p < 0.001, Holm–Bonferroni corrected), whereas formate and caproate did not differ significantly from blanks due to instrumental or solvent-related limitations (Fig. 1E). Despite modest inter-individual variability, all SCFA signals were consistently elevated in SC bees relative to MD and blank samples. While low levels of SCFAs in MD bees were associated with their gut microbial composition and bacterial loads (Supporting Information, Fig. S2, Table S1), we currently cannot explain the variation in SCFA between individuals in the colonized bees. A possible explanation would relate to eating behavior of individual bees altering the availability of substrates for bacterial fermentation.

Functional enrichment of the MS/MS-identified compounds revealed significant overrepresentation of metabolic pathways related to fatty acid metabolism, including butanoate, propanoate, and pyruvate metabolism, as well as glycolysis and fatty acid biosynthesis (Fig. 1F). Together, these results demonstrate that gut microbiota colonization profoundly reshapes the honey bee volatilome, primarily through modulation of fatty acid– associated metabolites that likely originate from microbial fermentation processes within the gut. This is in agreement with predictive functional analysis of the microbial communities found via 16S rRNA sequencing of the MD and SC guts (Fig. S4, Supporting Information).

### SESI-HRMS of live bees and LC-HRMS of bee guts cover distinct but complementary areas of the honey bee metabolome

To evaluate how headspace metabolomics by SESI-HRMS relates to conventional liquid chromatography-based metabolomics, we analyzed the water-extracted fraction of homogenated gut tissue of the same bees analyzed by SESI-HRMS (Fig. 1), using LC-HRMS (Fig. 2A). This paired analysis enabled a direct comparison of chemical coverage, compound annotation, and microbiota-associated metabolic shifts captured by each technique and body site.

**Fig. 2.**
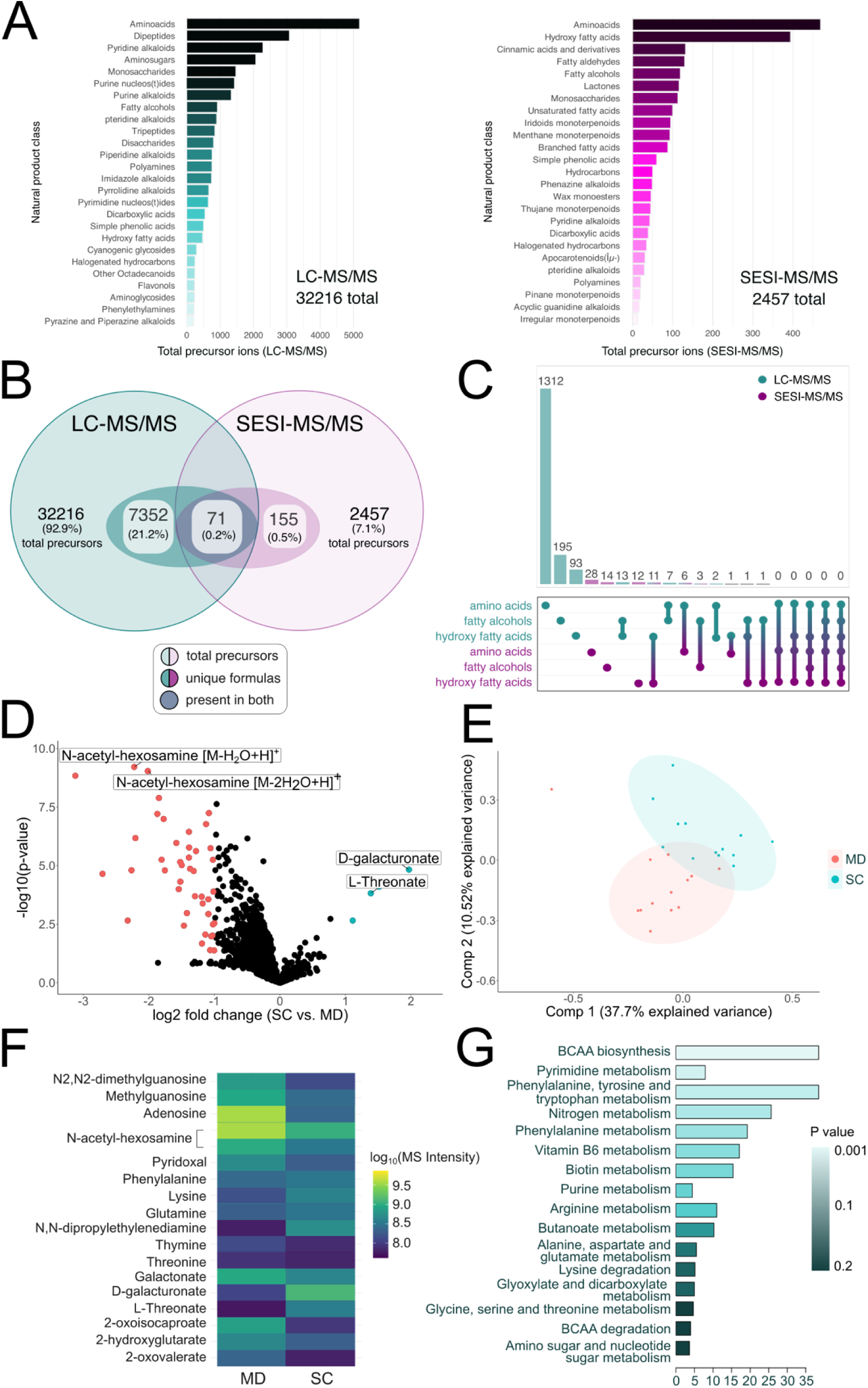
LC-HRMS comparison of the impact of gut microbiota on the metabolomic profiles of honey bee guts. **A)** Distribution of the most abundant natural product classes (NPCs) of the molecular formulas identified with LC-MS/MS (left) and SESI-MS/MS (right). **B)** Venn diagram showing the shared distribution of all the detected and putatively identified precursor ions in LC-MS/MS and SESI-MS/MS. Outside oval represents the total number of identified precursor ions, inside oval represents the non-repeating molecular formulas which could be assigned based on MS/MS spectra. **C)** Upset plot showing the overlap of compounds classified as amino acids, fatty alcohols, and hydroxy fatty acids between LC-MS/MS and SESI-MS/MS. **D)** Volcano plot showing the differential abundances of features found in MD and SC bees via LC-HRMS. Features with p < 0.05 and absolute fold change greater than 2 are highlighted in blue (higher in SC) and red (higher in MD). **E)** PLS-DA plot showing the distribution of samples belonging to the gut microbiota-depleted (MD) and synthetic community (SC) groups via LC-HRMS. **F)** Heatmap showing the logarithmic MS intensity of the most significantly different compounds in SC and MD bee guts by LC-HRMS. The intensities correspond to MS/MS-confirmed m/z values of identified metabolites. **G)** Enrichment analysis of the differentially abundant MS/MS-identified compounds from panel F (MetaboAnalyst 5.0, KEGG metabolite set).

Using the CANOPUS classification algorithm, we found that amino acids were the most abundant natural product class detected in both LC-MS/MS and SESI-MS/MS (Fig. 2A). However, the two platforms captured largely distinct molecular populations. LC-MS/MS detected a total of 32,216 precursor ions, representing 92.9 % of all features, whereas SESI-MS/MS detected 2,457 precursor ions (7.1 %). Only 71 molecular formulas (0.2 %) were shared between the two datasets (Fig. 2B). Among these, only six formulas corresponding to amino acids overlapped between the methods (Fig. 2C), confirming that SESI-MS/MS and LC-MS/MS probe separate regions of the honey bee metabolome.

The remaining natural product classes were largely associated with the phase of the sample. LC-MS/MS was dominated by amino acids, peptides, alkaloids and sugars, whereas the most abundant molecules in SESI-MS/MS constituted hydroxy fatty acids, cinnamic acids (plant-derived aromatic compounds present in honey, pollen, and bee propolis) and fatty alcohols and aldehydes (Supporting Information).

Overall, we found a much greater number of precursor ions in LC-MS/MS than SESI-MS/MS. Out of these, 7352 (21.2%) were attributable to unique molecular formulas, which resulted in many more hits being identifiable with MS/MS (Supporting Information).

Comparison of the shared m/z features found in the gut of MD and SC bees by LC-HRMS revealed pronounced group-specific metabolic differences (Fig. 2D-F). In total, most significantly altered metabolites were elevated in MD bees, paralleling the pattern observed in SESI-HRMS (Fig. 2D). PLS-DA demonstrated clear separation between MD and SC samples (Fig. 2E), indicating that gut microbial colonization strongly influences gut metabolite composition. Predominantly, the compounds upregulated in MD bees were nucleosides (guanosine, adenosine), amino sugars, and keto acids, which accumulate during incomplete breakdown of branched-chain amino acids (BCAAs). Additionally, we compared the intensities of several amino acids implicated in previous works on gnotobiotic honey bees, and found phenylalanine, glutamine, and lysine to be higher in SC, although these changes were modest relative to the more pronounced compounds (Fig. 2F).

The most discriminant compounds identified by MS/MS included nucleosides (adenosine, methylguanosine, N²,N²-dimethylguanosine), amino acids (phenylalanine, lysine, glutamine), and intermediates of branched-chain amino acid (BCAA) catabolism such as 2-oxo-isocaproate and 2-hydroxyglutarate (Fig. 2F). The most statistically significant feature, annotated as N-acetyl-hexosamine, appeared as two distinct ions differing by neutral water losses. While these m/z values could represent the same molecular species, it is metabolically more likely several amino sugar epimers (e.g., N-acetyl-glucosamine, N-acetyl-galactosamine), indistinguishable without targeted follow-up such as derivatization or selected high-resolution ion mobility.

To assess the metabolic impact of the differentially abundant metabolites, we applied the same enrichment analysis used for SESI-HRMS (Fig. 2G). LC-HRMS data revealed strong representation of BCAA biosynthesis and degradation, pyrimidine and purine metabolism, phenylalanine, tyrosine, and tryptophan metabolism, and vitamin B₆ metabolism, confirming the earlier findings (Fig. 2D, F). Consistent with the SESI-HRMS dataset, butanoate and glyoxylate/dicarboxylate metabolism also appeared significantly enriched, supporting the notion that gut microbiota modulate host amino acid and fatty acid biosynthesis pathways across both volatile and non-volatile metabolomic spaces.

Collectively, these results show that SESI-HRMS on live bees and LC-HRMS on isolated intestine provide complementary insights into the honey bee metabolome, with LC-HRMS capturing hydrophilic gut metabolites linked to amino sugar, amino acid and nucleotide metabolism, and SESI-HRMS profiling the corresponding volatile fermentation products and fatty acids emitted by the host.

### Butane-2,3-diol is a volatile metabolic signature of *S. marcescens* detectable in the headspace of infected honey bees

The Gram-negative species *S. marcescens* is an opportunistic pathogen of bees that causes elevated mortality in microbiota-depleted individuals but is largely suppressed in bees harboring a normal gut community.^11,38,39^ *Serratia* cultures are known to produce many volatile organic compounds, including various ketones, alcohols, and compounds involved in butanoate metabolism.^40–44^ We investigated the volatile signatures of *Serratia* by measuring isolated cultures grown under fully aerobic and anaerobic conditions, and then compared them to the headspace of honey bees colonized with either a defined synthetic community (SC), *S. marcescens* alone (SM), or both together (SCSM) (Fig. 3A). Each condition included four biological replicates with three bees per replicate (from four different hives), measured individually five days post-inoculation.

**Fig. 3.**
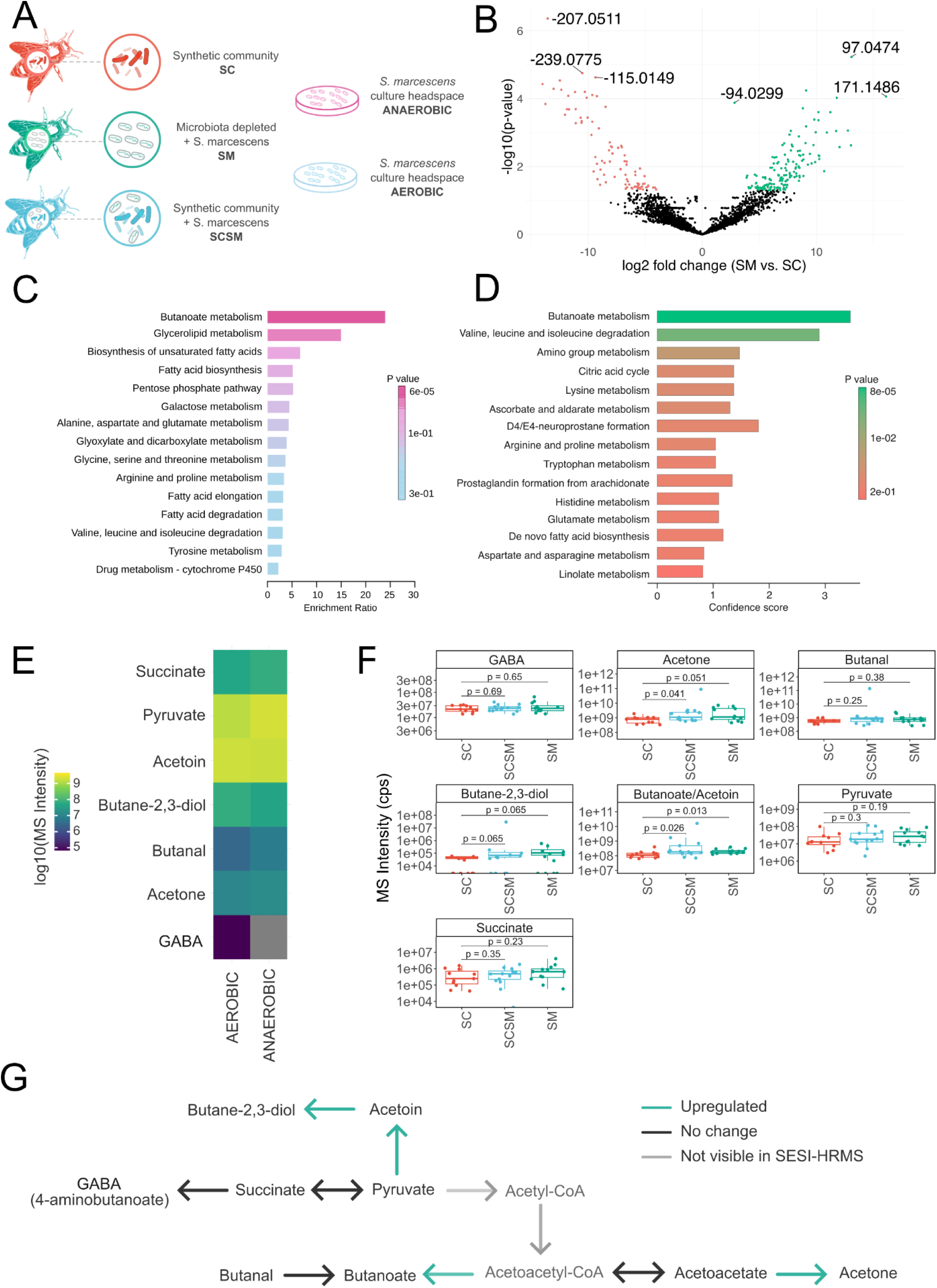
*S. marcescens* produces a distinct volatile signature detectable in the honey bee host. **A)** Schematic of the experimental design. SC – bees colonized with a synthetic community of bacteria. SM – bees monocolonized with *S. marcescens.* SCSM – bees co-colonized with *S. marcescens* and the synthetic community of bacteria. A total of four replicates were analyzed, with three bees per replicate. **B)** Volcano plot showing the differential abundances of features shared between the SC and SM groups. Features with p < 0.05 and fold change greater than 2 are highlighted in green (higher in SM) and red (higher in SC). **C)** Metabolic pathways enriched in the headspace of a liquid culture of an *S. marcescens* isolate. **D)** Metabolic pathways enriched in the headspace of honey bees colonized only with *S. marcescens.* **E)** Heatmap showing the relative abundance of m/z values found in the headspace of liquid cultures of *S. marcescens,* from the butanoate pathway (KEGG). The color of each tile corresponds to the logarithmic MS intensity of the m/z value corresponding to the labeled compound. **F)** Mean ion counts corresponding to compounds from the butanoate metabolism pathway of *Apis mellifera* (KEGG), matched by accurate mass. Statistical significance was measured by Welch’s t-test, adjusted for multiple comparisons using the Holm-Bonferroni method. P-value cutoffs: * - p < 0.05, ** - p < 0.01, *** - p < 0.001, **** - p < 0.0001. **G)** Proposed mechanism by which *Serratia* funnels the pyruvate surplus from the host into the production of butane-2,3-diol, butyrate (butanoate) and acetone visible in the host’s headspace. Green arrows show the path towards compounds with upregulated intensity. Compounds marked as light grey are non-volatile and not visible in SESI-HRMS.

Out of the total 2969 m/z features shared between the differently colonized bees, we found 153 to be upregulated in *Serratia*-colonized bees (SM), and 87 upregulated in synthetic community bees (SC, Figure 3B). Twenty-five features were uniquely elevated in *Serratia*-colonized bees, regardless of background microbiota (both SM and SCSM, Fig. S9). The subset of features abundant in the *Serratia*-only cohort was subsequently matched with the corresponding metabolic pathways using the Mummichog algorithm (MS1-based metabolic network prediction)^45^, assigning the highest number of features to butanoate metabolism and branch-chained amino acid metabolism (Fig. 3D).

To confirm a bacterial origin for these signals, we analyzed the headspace of *S. marcescens* grown in liquid culture. The strongest enrichment again corresponded to butanoate metabolism, followed by pathways linked to fatty acid biosynthesis and degradation (Fig. 3C). Serratia cultures produced a range of volatile metabolites derived from pyruvate, including acetoin, butane-2,3-diol, butanal, acetone, succinate, and γ-aminobutyrate (GABA), with differing abundances under aerobic and anaerobic growth (Fig. 3E). We also found clear differences in produced VOC content between *S. marcescens* grown aerobically and anaerobically (Supporting Information, Figure S6).

The genus *Serratia* is known to be a very efficient producer of acetoin and 2,3-butanediol.^46,47^ Both metabolites are derived from pyruvate and form parts of the butanoate pathway in *S. marcescens.*^48^ We examined the intensities of m/z values corresponding to several downstream metabolites of pyruvate, including butane-2,3-diol and acetoin, and found them all to be present at high intensities in the headspace of liquid cultures with SESI-HRMS (Fig. 3E). When investigating the same compounds in *Serratia*-colonized honey bees, we saw a corresponding increase in the intensities of m/z values corresponding to pyruvate, succinate, acetoin, butane-2,3-diol, and acetone, which exactly matched the liquid culture data (Fig. 3E,F).

We note that acetoin and butanoate (butyrate) are isobaric species that yield overlapping pseudomolecular ions, complicating quantitative interpretation. MS/MS spectra contained fragments diagnostic for both compounds (Supporting Information), but only butanoate can be synthesized by the host, whereas acetoin and butane-2,3-diol are typical microbial products. Increased signal at m/z 87.0451 ([M–H]⁻) in the headspace of infected bees therefore likely reflects Serratia-derived acetoin rather than host-origin butanoate.

Together, these results reveal that S. marcescens profoundly alters the honey bee volatilome by diverting host pyruvate flux toward the production of acetoin, butane-2,3-diol, butyrate, and acetone, metabolites that form a distinctive volatile signature of infection (Fig. 3G). These findings suggest a mechanistic link between microbial pyruvate metabolism and the emission of diagnostic volatiles in infected bees.

## DISCUSSION

This study demonstrates that volatile metabolite profiling using SESI-HRMS provides a sensitive and non-invasive approach for characterizing host-microbe metabolic interactions in the honey bee. By integrating single-bee volatilomics with LC-HRMS analysis of gut extracts, we uncovered distinct chemical signatures associated with (i) gut microbiota colonization, (ii) microbiota depletion, and (iii) bacterial infection.

Volatilomics offers a convenient and non-invasive sampling and analysis strategy, compared to traditional experimental interventions in metabolomics. It aims to minimize the necessary handling and stress incurred on the animal, and has been successfully used in metabolomic studies in many other model organisms, from bacteria to humans.^15,16,49,50^ In this work, we used the Western honey bee as a gnotobiotic model with a relatively well-defined and simple gut microbial composition to examine the metabolic differences detectable in their volatilome using SESI-HRMS. SESI-HRMS has proven particularly effective in diverse biological systems, from bacterial cultures to mammalian breath analysis, and here we demonstrate applicability to an insect host.

Our initial examination of the headspace of bees show that the gut microbiota profoundly alters the host volatilome (Fig.1). Bees colonized with a defined SC displayed distinct volatile signatures compared to MD bees, particularly within pathways associated with fatty acid biosynthesis and energy metabolism. Fatty acids play a number of very important roles in bee health, including changes in learning and cognition, energy metabolism, cellular membrane structure, environmental adaptation, and immune responses.^33,51–53^ Among the volatiles enriched in colonized bees, stearate and palmitate dominated the profile, both key precursors for lipid biosynthesis and royal jelly constituents^54,55^, and are the most abundant FAs in honey bees overall.^54,56^ Additionally, we found higher on average concentrations of SCFAs, such as acetate, propionate, and butyrate in SC bees, which are some of the main products of gut microbial activity, and also substrates for many of the host’s metabolic processes.^12,57^

Our complementary LC-HRMS analysis confirmed extensive metabolic remodeling associated with microbiota colonization. MD exhibited over 13 times as many significantly more abundant ions (3728) as SC bees (Figure 2D). We attribute this disparity to the accumulation of undigested food matter that would normally be broken down by the gut microbiota. Previous studies have reported that the pectin-degradation product galacturonate accumulates in the ileums of colonized bees.^34^ Pectin, a major component of the honey bee diet (pollen), along with other indigestible compounds such as cellulose and hemicellulose, is decomposed by the microbial community, particularly *Gilliamella apicola*.^22,58,59^ Consistent with this, we found galacturonate to be the most upregulated compound in SC bees, together with sugar acids and amino acids that serve as beneficial nutrients for the host (Figure 2D-G). While the MS/MS signature is in congruence with galacturonate, it has recently been suggested that the intensity previously attributed to this ion in mass spectrometry may in fact arise from the isomeric glucuronate, produced by the honey bee host.^60^ The authors observed increased abundance of the downstream metabolite of glucuronate, ascorbate, in hindguts of colonized bees as opposed to germ-free ones. Partly in agreement with this, we observed L-threonate, the degradation product of ascorbate, to be significantly upregulated in SC bees.

We also found a high abundance of nucleosides and nucleobases in the guts of MD bees, particularly methylated guanosine, adenosine, and thymine (Fig. 2F). Adenosine is associated with neurotransmission and stress responses in *Apis mellifera* and higher levels of adenosine receptors were found in bodies of bees infected with the common deformed wing virus, whereas lower levels were found in their brains.^61^ This suggests that the systemic response to the lack of beneficial gut community resembles the one caused by an infection. Thus, gut dysbiosis provokes stress and immune pathways resembling pathogenic challenge, a pattern conserved across insects and vertebrates.

When comparing the compound identification capabilities of SESI-MS/MS and LC-MS/MS we found a very high number of successful high-confidence MS/MS matches in LC-MS/MS (Fig. 2A). Feature annotation is one of the biggest challenges in untargeted *-omics* techniques, which makes preseparation and MS^n^ capabilities a major advantage in mass spectrometry-based workflows. Direct infusion (DI) techniques, while offering high throughput and real-time analysis capability, do not have this advantage. This is the likely reason for 10 times lower MS/MS identification rate in SESI-MS (Figure 2A-C), and a known problem in DI-MS.

While we found amino acids to be the most highly represented group in both LC-MS/MS and SESI-MS/MS, we found little overlap between the actual formulas and structures (Fig. 2C). Amino acids generally have extremely low vapor pressures and are not considered volatile (e.g., at 25°C glycine has a vapor pressure of only 1.73×10^-8^ kPa compared to 30.6 kPa for acetone, an analyte that is very easily ionized by SESI).^62,63^ Indeed, when manually examining the actual compound assignments (Supporting Information), it becomes apparent that the amino acids visible in SESI are rarely alpha amino acids: instead, it’s VOCs like γ-aminobutyric (GABA) and aminocaproic acid which contribute to this natural product assignment. However, because it is challenging to carry out LC-HRMS and SESI-HRMS on exactly the same samples due to the dilute nature of ambient air, we cannot conclusively link this discrepancy to the volatility of these compounds, but it remains the strongest working hypothesis for the observed differences.

As a final proof-of-concept for the applicability of volatile analysis to monitoring honey bee metabolic responses and health, we found bees infected with *S. marcescens* to have a different volatile profile than the healthy controls (Fig. 3B). When examining the headspace of a liquid *S. marcescens* culture (Fig. 3C) and the headspace of *Serratia*-infected bees otherwise depleted of gut microbiota (SM bees, Fig. 3D), we found the butanoate pathway to be upregulated in both. Considering they are lacking a healthy gut microbial community, the SM bees should be low in butyrate (butanoate) and compounds involved in the butanoate pathway, this suggests *Serratia’s* activity or change in the butanoate pathway as response to infection. The bacterial headspace measurements revealed the pyruvate to butane-2,3-diol pathway to be highly active, and this is consistent with previous literature on *S. marcescens.* When tracing these m/z values in honey bees, we did find their intensities to be higher in SCSM and SM bees. Given that honey bees lack the enzymatic machinery to produce butane-2,3-diol, its detection provides strong evidence for active bacterial metabolism within the host.

Butane-2,3-diol is moderately volatile (182 °C boiling point, 0.03 kPa under standard conditions) and poorly ionizable via SESI considering its low MS intensity. However, it does not have many molecular interferants, except for its own isomers. Other common compounds with this molecular formula are mainly industrial solvents, unlikely to be found in the beehive. This makes butane-2,3-diol an interesting candidate for a potential volatile biomarker, capable of early detection of infection by *S. marcescens*, even in the presence of native bacterial community. Further research is required into the sensitivity and specificity of this biomarker, especially considering its modest volatility, and to explore the potential of SESI-HRMS to detect and differentiate a range of economically relevant bee pathogens at early stages of infection.

Methodologically, combining SESI-HRMS with LC-HRMS provides a multidimensional view of host-microbe metabolism, spanning both volatile and non-volatile molecular spaces. Looking forward, the miniaturization of SESI interfaces and the development of portable or field-deployable VOC sensors could enable real-time, hive-side monitoring of honey bee health. Chemical markers such as acetoin or butane-2,3-diol could be integrated into biosensor systems to detect early signs of dysbiosis, nutritional stress, or infection, offering practical applications for pollinator management and conservation. More broadly, extending volatilomics to other insects and animal models may illuminate general principles of host–microbe metabolic communication and reveal conserved volatile mediators of symbiosis, immunity, and disease.

The limitations of this work include honey bees analyzed from one season only, as well as lack of controls altering the bee diet prior to analysis (bees were fed sugar-water, likely to be microbiome-neutral). The experimental setup in its current form lacks internal standards for normalization, as it does not involve any pressure-controlled flow-mixing chamber after the measurement tube (Fig. 4). The technique is also relatively low-throughput, especially when measuring one bee at a time (about 5 minutes per bee, including a preceding blank measurement). Its main application could therefore be identifying biomarkers to develop easy-to-use molecular sensors, more than real-time testing.

**Fig. 4.**
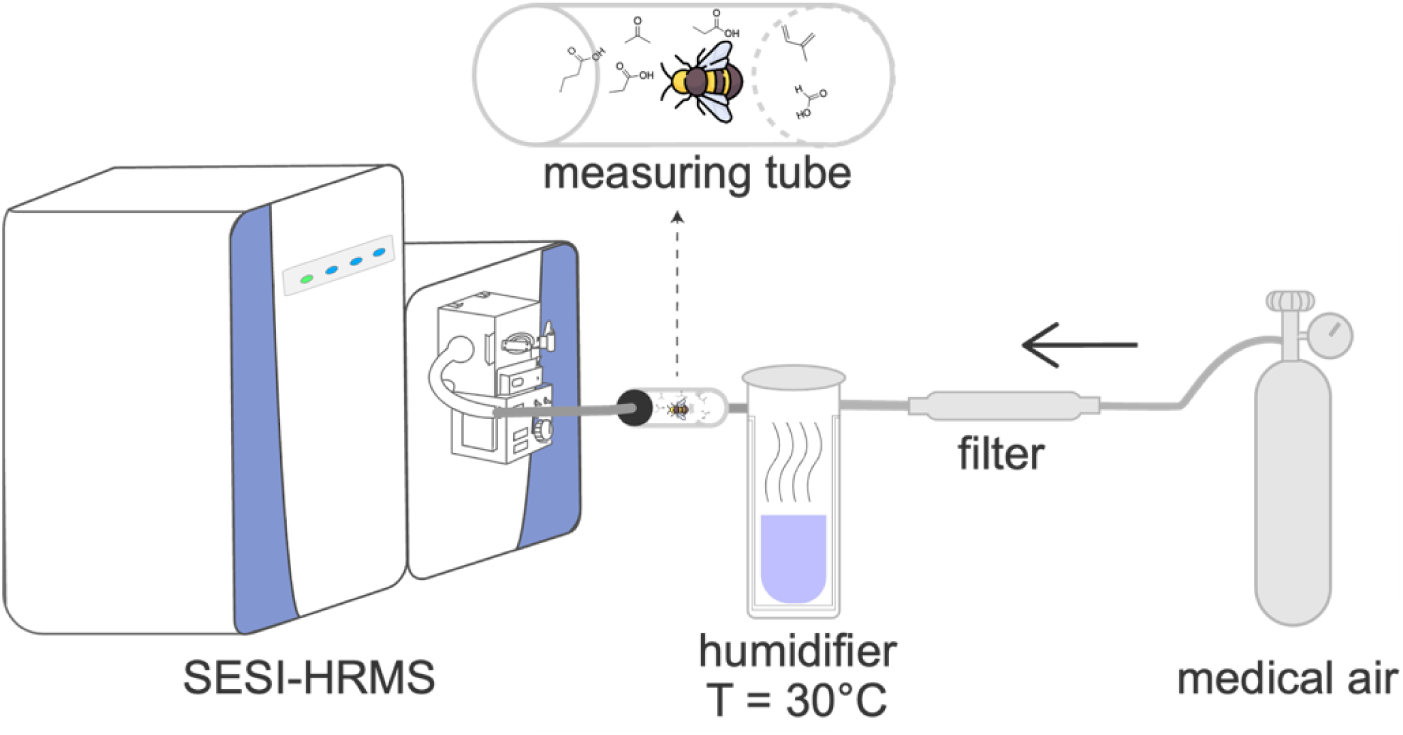
Schematic representation of the in-house headspace sampling system used for measuring volatiles released from honey bees.

## CONCLUSIONS

Volatile metabolites provide a real-time, non-invasive window into host-microbe metabolism, and the honey bee is an ideal system to exploit this view. By combining single-bee SESI-HRMS with LC-HRMS, we show that gut microbiota markedly reshape the bee volatilome, enriching fatty-acid-associated signals and short-chain fermentation products that typify a healthy, colonized state. LC-HRMS and SESI-HRMS proved complementary, capturing largely non-overlapping chemical spaces and together revealing novel differentiators between microbiota-depleted and colonized bees. Extending this framework to infection, we demonstrate that *S. marcescens* imposes a distinct volatile fingerprint consistent with rerouting host pyruvate through the butanoate pathway; notably, butane-2,3-diol emerges as a candidate biomarker of Serratia infection. These results establish volatilomics as a practical tool for monitoring bee health and microbiome function and lay the groundwork for field-deployable chemical sensing to detect dysbiosis or pathogen pressure in managed pollinators.

## MATERIALS AND METHODS

### Generation of Gnotobiotic Honey Bees

Late-stage pupae of *Apis mellifera*, identified by their pigmented eyes and light grey cuticle, were aseptically removed from brood frames and transferred to sterile emergence boxes. Pupae were incubated at 35 °C and 75% relative humidity for 3 days and supplied with sterile 1:1 (w/v) sucrose solution:PBS. Upon emergence, adult bees were randomly assigned to one of four experimental groups: microbiota-depleted (MD), colonized with a synthetic community of native gut bacteria (SC), monocolonized with *Serratia marcescens* (SM), or co-colonized with both the SC and *S. marcescens* (SCSM). All bacterial strains used in this study are listed in the Supporting Information. Strains were cultured under standard conditions, washed, and normalized to an optical density at 600 nm (OD₆₀₀) of 1.0. Aliquots were frozen in 25% glycerol at -80 °C until use. For colonization, bees were fed 5 μL of bacterial suspension diluted in phosphate-buffered saline (PBS) supplemented with 50% sucrose. SC group: Received a mix of 12 native gut strains at a combined at a final OD₆₀₀ of 0.1. SM group: Received a single strain of *S. marcescens* at OD₆₀₀ = 0.1. SCSM group: Received the SC mix (OD₆₀₀ = 0.1) supplemented with *S. marcescens* (OD₆₀₀ = 0.1). MD group: Received a mock inoculum containing only the PBS:sucrose vehicle. Bees were maintained under sterile conditions throughout the experiment and provided with autoclaved 1:1 sucrose solution. Experiments were performed 5 days post-inoculation. After SESI-MS measurements bees were immediately stored at -80 °C until further analysis. To minimize potential hive-specific effects, all experiments were biologically replicated using bees from four independent hives (n = 4). Where indicated, two technical replicates per treatment were also conducted using separate cages to account for cage-related variation.

### Real-time SESI-HRMS measurements of honey bee headspace

A set of air-tight transparent measurement tubes (2 cm diameter, 7 cm length, PMMA) was manufactured with Swagelok fittings and used as part of the in-house headspace sampling system (Figure 4). Just before the experiment, the bees were first stunned using dry ice and then transferred to their measuring tube. 0.3 L×min^-1^of humidified medical air (Pangas, Switzerland) was flushed through a carbon filter and into the bee tunnel, and subsequently introduced into the ion source. The humidifier, the bee tunnel and all gas tubing in between were kept at 30 ± 2 °C. After the measurements, bees were anesthetized with carbon dioxide and decapitated. The guts were extracted for LC-HRMS analysis and DNA extraction within 30 minutes of headspace measurements.

### Real-time SESI-HRMS measurements of *S. marcescens* headspace

*S. marcescens* isolates were grown on BHI-S plates and single colonies were then chosen as replicates for growth in liquid Sf-900 insect cell culture media. The cultures were grown either under aerobic or under fully anaerobic conditions, with three replicates per condition. The replicates were sampled during exponential growth (OD_600_ = 0.3) and transferred into sealed round-bottom flasks. To sample the headspace, 0.3 L×min^-1^ of humidified carrier gas (9:1 mixture of nitrogen to carbon dioxide, Pangas, Switzerland) was flushed through the round-bottom flask containing 3 mL of the bacterial culture, and introduced into the SESI source. The humidifier, the headspace sampler, and all gas tubing in between were kept at 30 ± 2 °C.

### Data acquisition by SESI-HRMS

Data was acquired using a commercial SESI source (Fossil Ion Tech, Spain) coupled to a Q Exactive Plus Orbitrap high-resolution mass spectrometer (ThermoFisher Scientific, Germany). During measurements, a pressure of ca. 8 x 10^-11^ mbar was maintained. The SESI sampling line and the ionization chamber were heated at 130 °C and 90 °C, respectively. For each measurement, both positive and negative ionization modes were recorded separately. The electrospray was generated using a 0.1% aqueous (LC-MS grade water, ThermoFisher Scientific) formic acid solution (99.5% formic acid, VWR Chemicals, Switzerland), passed through an electrospray capillary (outer diameter = 365 μm, inner diameter = 20 μm) under a positive pressure of 800 mbar. The sheath gas was kept at 15 a.u., and the auxiliary gas at 2 a.u. The electrospray voltage was set to ± 3.5 kV, depending on the polarity mode. The MS inlet capillary was heated to 250 °C. Automatic gain control in the C-trap was set to 1e6 and the maximum injection time to 500 ms. The mass resolution was kept at 140,000 for both full-MS and MS/MS scans, and the S-lens RF level was set to 50. For each polarity mode, MS1 spectra were recorded in profile mode, followed by 10 consecutive data-dependent MS2 scans in centroid mode.

32 individual honey bees were analyzed, wherein a single bee was measured at a time. Once proper air flow, airtightness, and heating were ensured, the measurement was started for a total of 4 minutes: 60 seconds for a full MS1 acquisition in positive ion mode, another 60 seconds for the negative ion mode, 60 seconds for DDA MS/MS measurements in positive ion mode, and 60 seconds in the negative ion mode. Additionally, blank measurements of the background ions present in humidified medical air passed through the measurement tube were taken before every replicate.

### Metabolite extraction for LC-HRMS

After excision, the honey bee gut samples were transferred into tubes containing zirconia beads (0.1 mm dia. Zirconia/Silica beads, Carl Roth, Switzerland). The wet weight of each sample was taken, and 10 times v/w Optima LC-MS grade water (FisherScientific, Switzerland) was added. The samples were homogenized using a Retsch MM 400 homogenizer at 30 Hz for 30 seconds. The samples were then split into two fractions: for metabolomics by and for DNA extraction. The metabolomics aliquots of 100 µl were further diluted 1:1 with Optima water. Following a previously established protocol^1^ the samples were incubated at 80°C and 1,400 rpm for 3 minutes, additionally vortexing the samples after each minute. This procedure was then followed by 5 minutes of centrifugation at 20,000 x g at 4°C, after which 150 µl of the supernatant was collected and centrifuged under the same conditions for another 30 minutes. Finally, the samples were again 10x diluted in LC-MS grade water and stored at -80°C.

### 16S rDNA sequencing of the gut bacterial loads

The homogenized samples for DNA extraction were processed using phenol-chloroform extraction and sodium acetate precipitation as described previously.^28^ The DNA quality was assessed using the 4200 TapeStation System (Agilent Technologies, Switzerland). Full-length amplicon metagenomic sequencing of the 16S rDNA was performed using the PacBio Revio platform (Novogene, Germany). Universal primers were designed using the sequence in conserved regions flanking at the upstream and downstream of the targeted hypervariable regions. The PacBio BAM file was split according to barcode and filtered to get clean data. The filtered data were analyzed by generating Amplicon Sequence Variants (ASVs) for species annotation. Subsequently, ASVs were used for taxonomic assignment and abundance distribution estimation. Prediction of metagenomic functions was performed using the PICRUSt2 algorithm.

### Data acquisition by LC-HRMS/MS

Samples were injected into a Vanquish Flex (ThermoFisher Scientific, Germany) liquid chromatograph coupled to a heated electrospray ion source (H-ESI) of an Orbitrap Exploris 240 mass spectrometer (ThermoFisher Scientific, Germany). Hydrophilic interaction chromatography was performed on a Waters Acquity BEH HILIC Column (1.7 µm, 2.1 mm x 100 mm). The mobile phase was a gradient mixture of ammonium formate (2 mM, aqueous), formic acid (0.1% v/v, pH 3), and acetonitrile. Detailed information on the gradient can be found in the Supporting Information. Absorbance data for each run were collected at 254 nm and 280 nm. The mass spectrometer was operated in both the positive and negative ionization modes, each recorded in a separate run. In both modes, the sheath gas was set to 30 a.u., the auxiliary nebulizer gas to 10 a.u. at 400 °C, and the ion transfer tube was heated to 350 °C. The S-lens RF voltage was kept at 0. In positive mode, the capillary voltage was set to 3.4 kV, and in negative mode to 2.5 kV. Data acquisition was performed with full-MS (50-750 m/z) scans recorded in profile mode, followed by a series of twenty consecutive data-dependent MS/MS (ddMS^2^) scans recorded in centroid mode, with an intensity threshold and a sixty-second dynamic exclusion window filter. The ddMS^2^ scans were acquired with stepped normalized collision energies of 5, 30, 70, 150, and 200. In both full-MS and MS/MS scan modes, the mass spectrometer resolution was set to 120,000 with a standard (1e6) AGC target and automatic injection time adjustment.

### Data processing

Raw MS data files in the proprietary ThermoFisher Scientific RAW file format were first converted into the open mzML format using MSConvert (ProteoWizard package).^64^ Raw LC chromatograms were directly exported from the Chromeleon Chromatography Data System in the TXT file format. The mzML and TXT files were subsequently analyzed using LCMSpector, a standalone user-interface application for viewing and analyzing LC-MS data.^65^ The processed CSV files, containing all the targeted primary ions, their absorbance values and MS intensities, as well as concentrations extrapolated from calibration curves of standard samples, were then exported and further analyzed with R. All the data visualization and post processing scripts can be found in the Supporting Information.

Data-dependent acquisition tandem mass spectra (DDA-MS2) were processed and annotated with SIRIUS.^66^ Detailed information containing the putative assignments of fragment ions can be found in the Supporting Information.

## AUTHOR CONTRIBUTIONS

S.M.G. and M.F. together with J.L. designed the study. E.S. and P.E. supervised the study. S.M.G. and A.P. cultivated, colonized, and transported the honey bees. S.M.G., A.P., and M.F. performed the SESI-HRMS measurements. M.F. and L.Z. performed the LC-HRMS measurements. J.L. designed the initial headspace sampling system, conducted preliminary measurements, and was consulted on the analysis process. E.C.B. supervised and took part in the processing of metabolomic samples and DNA extraction. M.F. processed and analyzed the SESI-HRMS and LC-HRMS data. R.Z., P.E., and E.S. critically read the first draft of the manuscript. The manuscript was written through the contributions of all authors. All authors have edited the manuscript and given approval for the final version.

## ACKNOWLEDGMENTS

We thank Laurence Cuche of the Building Directorate Office of Spatial Development of the Canton of Zurich for providing wild honey bees for additional experimentation. We also thank the Departmental Mechanical Workshop for manufacturing and providing the measurement tubes and other parts of the bee headspace analysis system. We thank the *FBM décanat* for the support provided during the parental leave of S.M.G. This work was funded by the NCCR Microbiomes, a National Centre of Competence in Research, funded by the Swiss National Science Foundation (grant no. 180575) and a Swiss National Science Foundation project grant (grant no. 179487) awarded to P.E.

## DATA AVAILABILITY STATEMENT

The original data used in this publication will be made available in a curated data archive at ETH Zürich (https://www.research-collection.ethz.ch).

